# Fluorescence-barcoded cell lines stably expressing membrane-anchored influenza neuraminidases

**DOI:** 10.1101/2025.01.01.631020

**Authors:** Joel Finney, Masayuki Kuraoka, Shengli Song, Akiko Watanabe, Xiaoe Liang, Dongmei Liao, M. Anthony Moody, Emmanuel B. Walter, Stephen C. Harrison, Garnett Kelsoe

## Abstract

The discovery of broadly protective antibodies to the influenza virus neuraminidase (NA) has raised interest in NA as a vaccine target. However, recombinant, solubilized tetrameric NA ectodomains are often challenging to express and isolate, hindering the study of anti-NA humoral responses. To address this obstacle, we established a panel of 22 non-adherent cell lines stably expressing native, historical N1, N2, N3, N9, and NB NAs anchored on the cell surface. The cell lines are barcoded with fluorescent proteins, enabling high-throughput, 16-plex analyses of antibody binding with commonly available flow cytometers. The cell lines were at least as efficient as a Luminex multiplex binding assay at identifying NA antibodies from a library of unselected clonal IgGs derived from human memory B cells. The cell lines were also useful for measuring the magnitude and breadth of the serum antibody response elicited by experimental infection of rhesus macaques with influenza virus. The membrane-anchored NAs are catalytically active and are compatible with established sialidase activity assays. NA-expressing K530 cell lines therefore represent a useful tool for studying NA immunity and evaluating influenza vaccine efficacy.

## INTRODUCTION

Influenza A and B viruses (IAVs and IBVs) are substantial public health burdens. Two A subtypes (H1N1 and H3N2) and one B lineage (Victoria) currently circulate among humans (1). Another B lineage (Yamagata) that recently circulated may have gone extinct (2, 3). Additional flu strains circulating in animal populations can also cause zoonosis (4, 5). Seasonal flu vaccines offer important protection against disease, but they elicit narrow immunity against the proteins of the viral strains included in the vaccine, necessitating annual vaccine updates as pressure from herd immunity selects for escape mutations (6–9). Thus, the goal of many next-generation flu vaccines is to elicit antibody (Ab)-mediated protection against serologically divergent viral strains (10, 11). Much of the flu vaccine research of the last 15 years has focused on Ab responses to hemagglutinin (HA), the major glycoprotein on the virion surface, which mediates attachment to sialylated cellular receptors and catalyzes membrane fusion to release the viral genome into the host cell. The viral neuraminidase (NA), a less abundant glycoprotein whose function is to cleave sialic acids from cellular receptors to enable viral egress, is also an immune system target (12). The recent discovery of protective Abs that recognize conserved epitopes on NAs from divergent IAVs and IBVs has renewed interest in designing vaccines that target NA (13–18).

Recombinant, full-length, solubilized NA ectodomains are frequently challenging to express, impeding study of NA immunity. The typical expression construct for recombinant NA comprises only the globular head domain (i.e., excluding the N-terminal stalk region)(18–20), and the yield of the resultant protein is 10- to 100-fold lower than that routinely achieved for recombinant HA ectodomains (20). Full-length NAs have been expressed in the native, membrane-anchored form on enveloped virus-like particles (14, 15), but expression and purification of virus-like particles requires access to specialized equipment, which may be a barrier for some laboratories. Others have expressed full-length NAs on transiently transfected, mammalian cell lines (14, 21), but these transfected cell lines are neither practical for multiplexed analysis of Ab binding breadth nor easily shared between laboratories.

We have used panels of fluorescence-barcoded K530 cell lines stably expressing membrane-anchored HAs or major histocompatibility complex (MHC) proteins to determine the binding breadth and avidity of relevant Abs in a flow cytometry assay (22–24). These non-adherent cell lines are easy to culture, have short doubling times, and can be used to study Ab binding to up to 16 different antigens simultaneously (23). The barcodes and fluorescence intensity of Ab binding can be detected using 405 nm, 488 nm, and 633 nm excitation lasers and standard emission filter sets available on most flow cytometers, with good inter-assay reproducibility (23). Here, we have adapted the barcoded cell lines to express native NAs from historical isolates of influenza A and B viruses that have circulated in human or animal populations.

## RESULTS

### Standard NA monoclonal Abs brightly and specifically label K530-NA cell lines

We selected monoclonal, fluorescently barcoded K530 cell lines (23) stably expressing native, full-length NAs representing 22 historical isolates of IAV or IBV (Table I, Supplemental Table I). Another barcoded cell line expressing no NA controls for binding specificity. Up to 16 cell lines can be pooled for a single assay, with two different NA options for some of the barcodes. Collectively, the cell lines express NAs from the N1, N2, N3, and N9 subtypes of IAV, including N1s spanning 44 years (1977–2021) of human isolates and N2s spanning 65 years (1957–2021) of human isolates. NAs representing the Victoria and Yamagata lineages of IBV are also included.

**Table 1.** Fluorescence-barcoded cell lines expressing recombinant, membrane-anchored NAs.

|  | Option 1 |  | Option 2 |  |
| --- | --- | --- | --- | --- |
| Fluorescent barcode<sup>a</sup> | Source of NA | Abbreviated NA name | Source of NA | Abbreviated NA name |
| 0000 | – | – | – | – |
| 0001 | A/Solomon Islands/03/2006 (H1N1) | N1.SI06 | A/Vietnam/CL01/2004 (H5N1) | N1.VN04 |
| 0010 | A/Sydney/5/2021 (H1N1) | N1.SYD21 | – | – |
| 0011 | A/Hong Kong/4801/2014 (H3N2) | N2.HK14 | – | – |
| 0100 | A/Hong Kong/1/1968 (H3N2) | N2.HK68 | – | – |
| 0101 | A/USSR/90/1977 (H1N1) | N1.USSR77 | A/Japan/305/1957 (H2N2) | N2.JP57 |
| 0110 | A/Michigan/45/2015 (H1N1) | N1.MI15 | A/Hong Kong/1144/1999 (H3N2) | N2.HK99 |
| 0111 | A/Bilthoven/1761/1976 (H3N2) | N2.BH76 | A/Darwin/9/2021 (H3N2) | N2.DW21 |
| 1000 | A/Kansas/14/2017 (H3N2) | N2.KS17 | A/Jiangsu/428/2021 (H10N3) | N3.JS21 |
| 1001 | A/Memphis/4/1987 (H1N1) | N1.ME87 | – | – |
| 1010 | B/Brisbane/60/2008 (Victoria) | NB.BN08 | – | – |
| 1011 | A/California/07/2009 (H1N1) | N1.CA09 | – | – |
| 1100 | A/Brisbane/8/1996 (H3N2) | N2.BN96 | A/Shanghai/02/2013 (H7N9) | N9.SH13 |
| 1101 | A/Beijing/353/1989 (H3N2) | N2.BJ89 | – | – |
| 1110 | A/Perth/16/2009 (H3N2) | N2.PE09 | A/tern/Australia/G70C/1975 (H11N9) | N9.AU75 |
| 1111 | B/Phuket/3073/2013 (Yamagata) | NB.PK13 | – | – |

Notes:
<sup>a</sup> Expression of a fluorescent protein is indicated with a “1”, whereas “0” indicates the protein is not expressed. The “ones” place (0001) denotes expression of eBFP2, the “tens” place (0010) denotes mTurquoise, the “hundreds” place (0100) denotes mNeonGreen, and the “thousands” place (1000) denotes mCardinal.

In a flow cytometry assay, recombinant IgGs (rIgGs) representing well-characterized, monoclonal Abs (mAbs) against the NA catalytic site (13–15, 18) brightly and specifically labeled the K530-NA cells, but not control cells that expressed no NA (Fig. 1). As reported, 1G01 (15, 18), FNI17 (14) and DA03E17 (15) bound NAs of diverse subtypes from phylogenetic groups 1 and 2 of IAV (Fig. 1). FNI17 and DA03E17 also bound NAs from both lineages of IBV, as described (14, 15). Although 1G01 was reported to recognize NAs from some strains of IBV, 1G01 binds IBV NAs much more weakly than IAV NAs, and even 50 μg/ml of 1G01 cannot neutralize B/Brisbane/60/2008 or B/Phuket/3073/2013 virus in vitro (14, 15, 18). Thus, our observation that 1G01 did not label K530 cells expressing NB.BN08 or NB.PK13 (Fig. 1) agrees with prior reports. In contrast, as expected, 1G05 mAb (13) avidly bound NAs from both IBV lineages, but did not bind IAV NAs (Fig. 1). S1V2-72, a control IgG that specifically binds influenza hemagglutinin (22), did not bind K530-NA cells.

**FIG 1.**
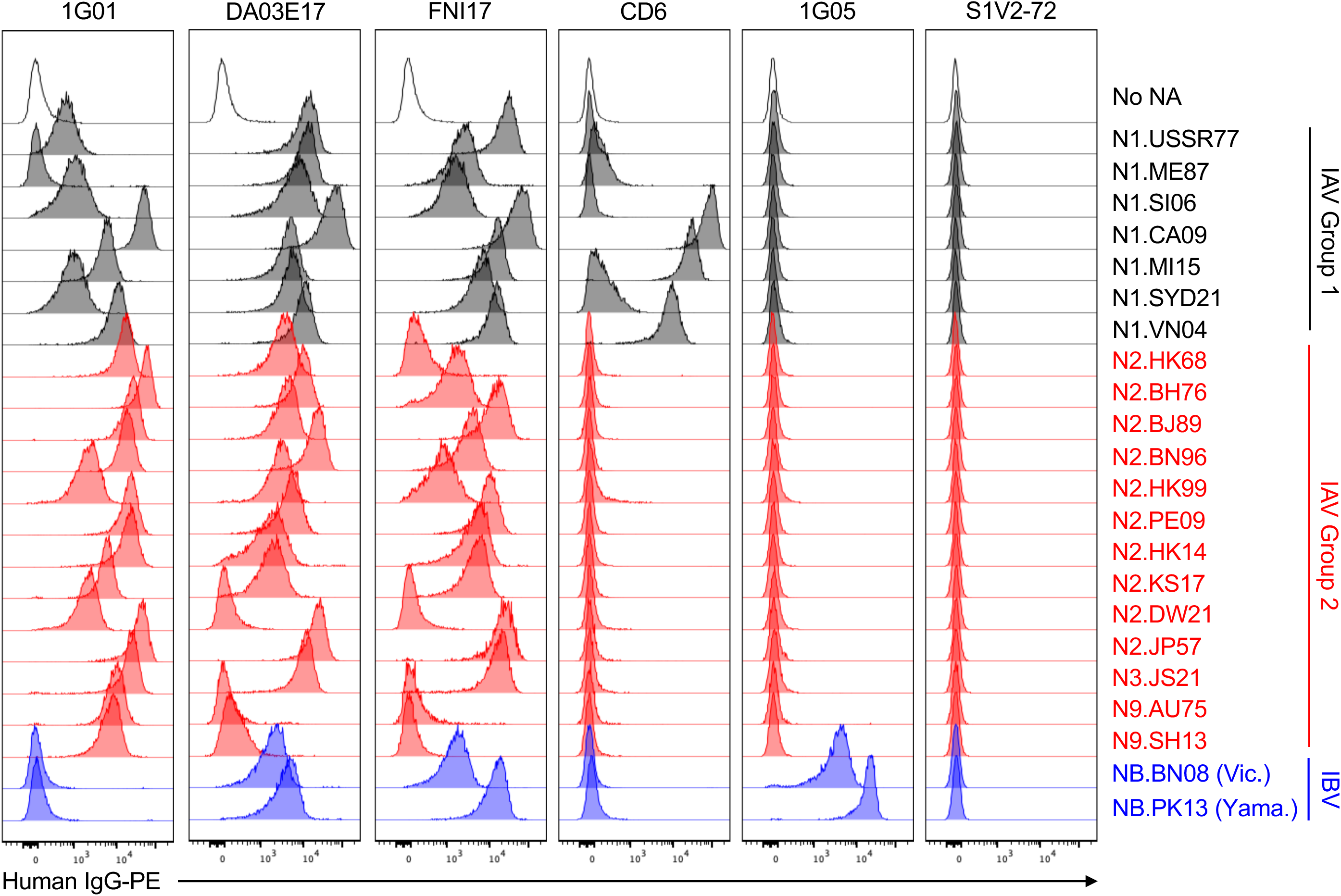
Standard NA mAbs brightly and specifically label K530-NA cell lines. Shown are flow cytometry histograms depicting the binding of recombinant IgG versions of standard NA mAbs to K530 cell lines expressing membrane-anchored NAs. Each row corresponds to a monoclonal cell line stably expressing a single type of NA. Each column corresponds to the binding profile for a single mAb. MAbs were incubated with pooled cell lines comprising Option 1 or Option 2, and the resultant data were concatenated into a single figure.

Standard mAb CD6 was reported to bind N1.CA09 with high affinity (21). We observed tight binding of CD6 to N1.CA09-expressing K530 cells (Fig. 1). We also discovered that CD6 avidly bound N1.MI15, a descendant of N1.CA09, and weakly bound a later descendant, N1.SYD21. CD6 also recognized N1.VN04, which was not detected in an earlier report using glutaraldehyde-fixed, transiently transfected 293T cells expressing membrane-anchored N1.VN04 (21). However, our observation that CD6 binds N1.VN04 is consistent with its modest neutralizing activity toward A/Vietnam/1203/2004 (H5N1) virus (21). Because the CD6 epitope evenly spans adjacent protomers of the NA homotetramer (21), bright labeling of NA-expressing K530 cells by CD6 implies that the NA is in its native, multimeric form.

### K530-NA cells perform as well as a Luminex binding assay for high-throughput discovery of NA Abs

To test whether K530-NA cells can be used to identify new NA Abs, we sorted individual memory B (Bmem) cells from peripheral blood mononuclear cells of a healthy, teenaged donor (T3) 14 days after immunization with the 2019-2020 seasonal influenza vaccine (for an example of the sorting strategy, see Supp. Fig 1A) (25). During sorting, Bmem cells were not selected for binding to any antigen. From cultures of these Bmem cells, we obtained 1,847 clonal IgG-containing culture supernatants, which we screened for NA-binding activity both by Luminex assay (using tetrameric NA head constructs representing N1s, N2s, and NBs) and by multiplex flow cytometry (using the nine K530-NA cell lines we had generated at the time of this experiment). The Luminex screen identified five clonal IgGs that avidly bound NA: T3-P4A8 and T3-P10F5, which bound N1s from new pandemic H1N1 strains; T3-P17F8 and T3-P27B8, which bound N2.HK14; and T3-P15C5, which bound NBs from both IBV lineages (Fig. 2A). The Luminex screen also identified 11 samples with weak or borderline binding to NA. The flow cytometry screen with K530-NA cells (Fig. 2B) also identified the same five IgGs with avid NA binding, plus two IgGs with modest NA binding: T3-P20D5, which cross-reacted with numerous N1s, N2s, and NB; and T3-P28D3, which bound N2.BJ89, an NA not included in the Luminex screen. A third IgG, T3-P35E4, cross-reacted with N1s, N2s, and NB, but also weakly bound K530 cells that did not express NA; this sample may be auto- or polyreactive. The K530-NA cells also identified several IgGs with weak/borderline NA binding. Thus, for high-throughput identification of NA Abs, K530-NA cells perform at least as well as a Luminex assay using soluble NA heads; the K530 cells also have the advantage of potentially identifying Abs to the NA stalk domain.

**FIG. 2.**
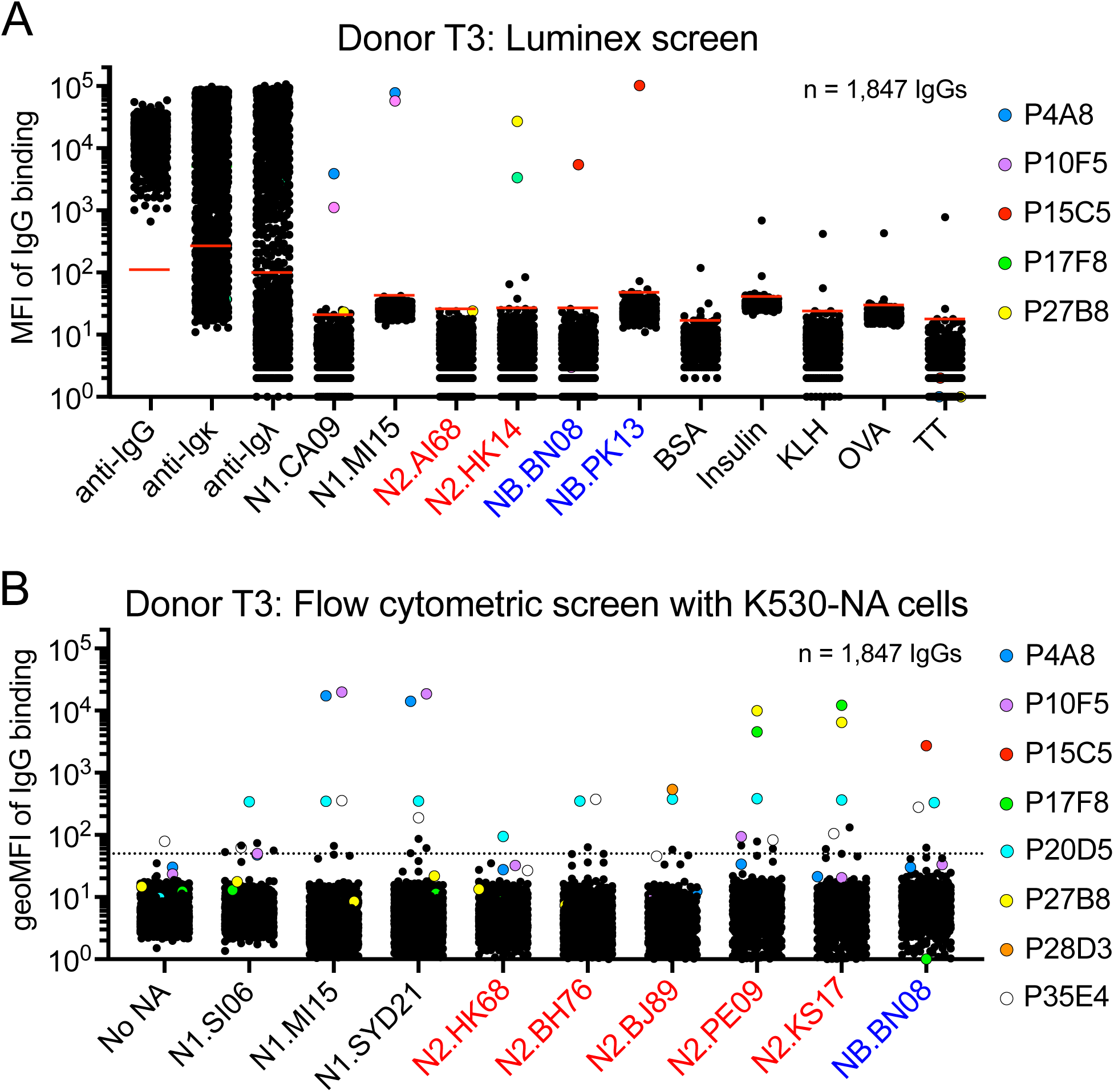
K530-NA cell lines perform as well as a Luminex assay for identifying NA mAbs. A) Luminex assay results showing the median fluorescence intensity (MFI) of antigen binding for clonal IgG-containing culture supernatants from Bmem isolated from donor T3. Bovine serum albumin (BSA), insulin, keyhole limpet hemocyanin (KLH), chicken ovalbumin (OVA), and tetanus toxoid (TT) serve as binding-specificity controls. Short, red, horizontal lines in each column denote the limit of detection (LOD), which was calculated as six standard deviations above the mean signal produced by cultures containing no B cells. B) Results of a high-throughput, flow cytometry-based screen using K530-NA cell lines. Each symbol depicts the geometric mean fluorescence intensity (geoMFI) of clonal IgG binding to a K530-NA cell line. The dashed, horizontal line denotes the threshold for binding. Each symbol in (A) and (B) represents a single, clonal IgG-containing culture supernatant. Clonal IgGs of interest are denoted with uniquely colored symbols.

We expressed rIgGs representing NA mAbs identified from donor T3 and two other teenaged donors (Supp. Fig. 1B-1C), and tested the binding breadth of these rIgGs using all 22 K530-NA cell lines (Fig. 3). The rIgGs recapitulated the binding profiles observed for the corresponding culture supernatant IgGs in the initial Luminex or K530-based screens (Fig. 3). Several of the rIgGs also bound additional NAs not included in the initial screens. Each rIgG bound only one NA subtype, except T3-P20D5, which bound various N1s, N2s, N3.JS21, and NB.BN08. T3-P20D5 bound these NAs weakly, but specifically: when we set the sensitivity of the flow cytometer’s phycoerythrin (PE) detector to a level that caused the signal from higher-affinity standard mAbs to approach or exceed the detector limit, T3-P20D5 clearly labeled some K530-NA cell lines, but not others (Supp. Fig 2A). It is unclear why T3-P20D5 did not bind the tetrameric NA heads used in the Luminex screen (Supp. Fig. 2B); one possibility is that T3-P20D5 recognizes an epitope in the NA stalk domain.

**FIG. 3.**
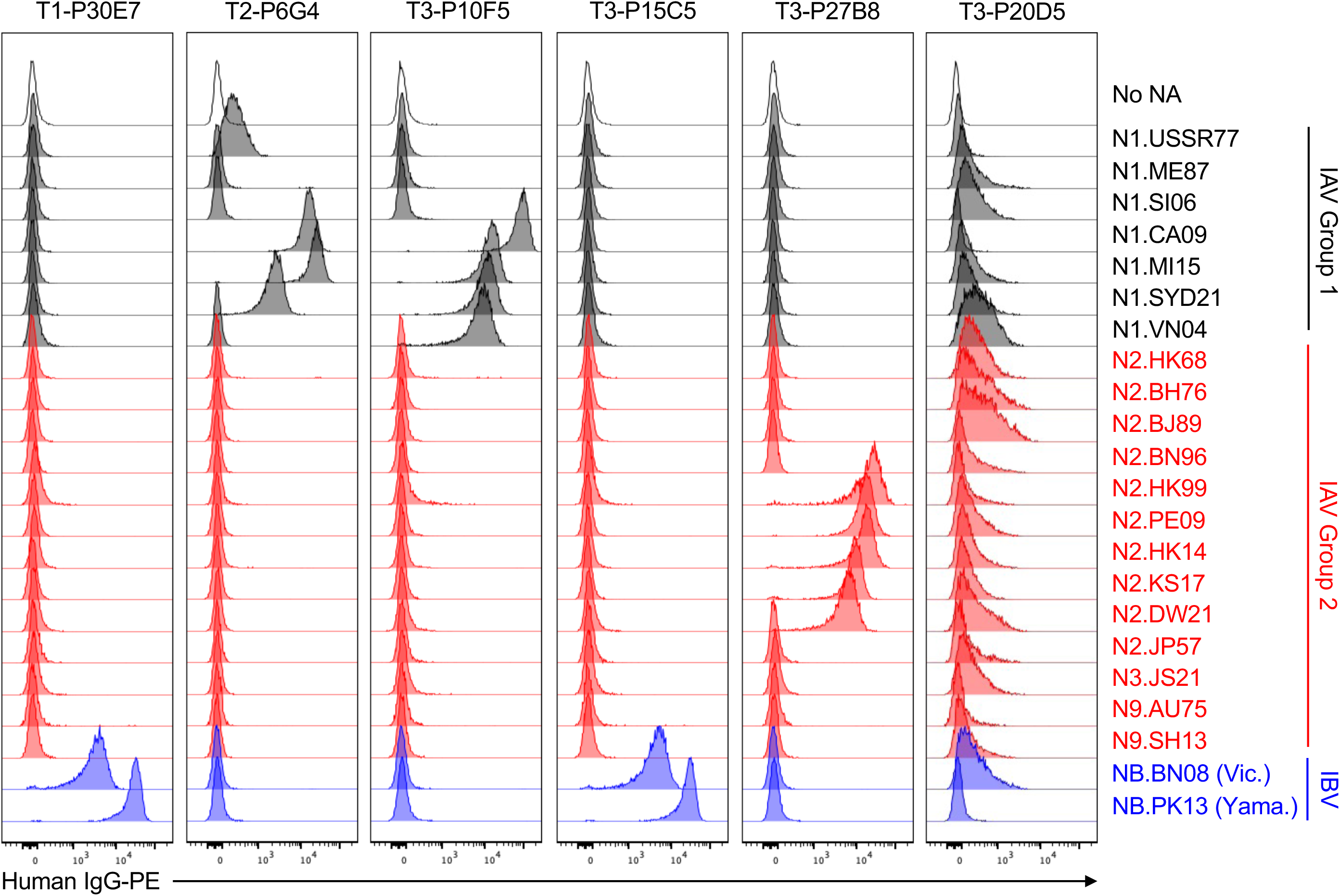
Use of K530-NA cells to determine the binding breadth of newly identified NA mAbs. Shown are flow cytometry histograms depicting the binding of recombinant IgG versions of NA mAbs from donors T1, T2, or T3 to K530 cell lines expressing membrane-anchored NAs, as in Fig. 1.

### K530-NA cells detect anti-NA serum Abs elicited by influenza infection

We used K530-NA cell lines to analyze the serum Ab response to IAV infection in rhesus macaques (*Macaca mulatta*) (26). At the start of the experiment, each monkey was seronegative for influenza virus exposure, as determined by hemagglutination inhibition titer (data not shown). We infected three animals with A/Aichi/2/1968 (H3N2) influenza virus, and immunized two control animals with recombinant influenza HA (26). Diluted pre-immune or immune plasma from each animal (Supplemental Fig. 3A) was incubated with pooled K530-NA cell lines, and then the fluorescence intensity of IgG labeling was detected by flow cytometry.

As expected, vaccination with recombinant HA did not elicit NA-binding serum IgG (Supplemental Fig. 3B). In contrast, infection with H3N2 influenza virus elicited N2-specific serum IgG responses (Fig. 4). Using the fold-change of the intensity of IgG labeling as an indirect measure of the quantity of N2-specific antibody elicited by infection or immunization, we observed that infection-elicited IgG primarily bound autologous antigen (N2.HK68), but also reacted substantially with N2.JP57, a representative of the 1957 H2N2 pandemic virus that preceded the emergence of H3N2 in 1968 (Fig. 4B). Infection-elicited IgG cross-reacted modestly with later isolates of N2, but did not bind N2s from 2009 or after. The elicited IgG did not cross-react with non-N2 subtypes, for although each animal’s serum contained some N1-, N3-, N9- and/or NB-reactive IgG before infection, the quantities of these IgGs did not increase after IAV exposure (Fig. 4 and Supplemental Fig. 3C). Thus, K530-NA cell lines are useful for analyzing serum Abs as well as mAbs.

**FIG. 4.**
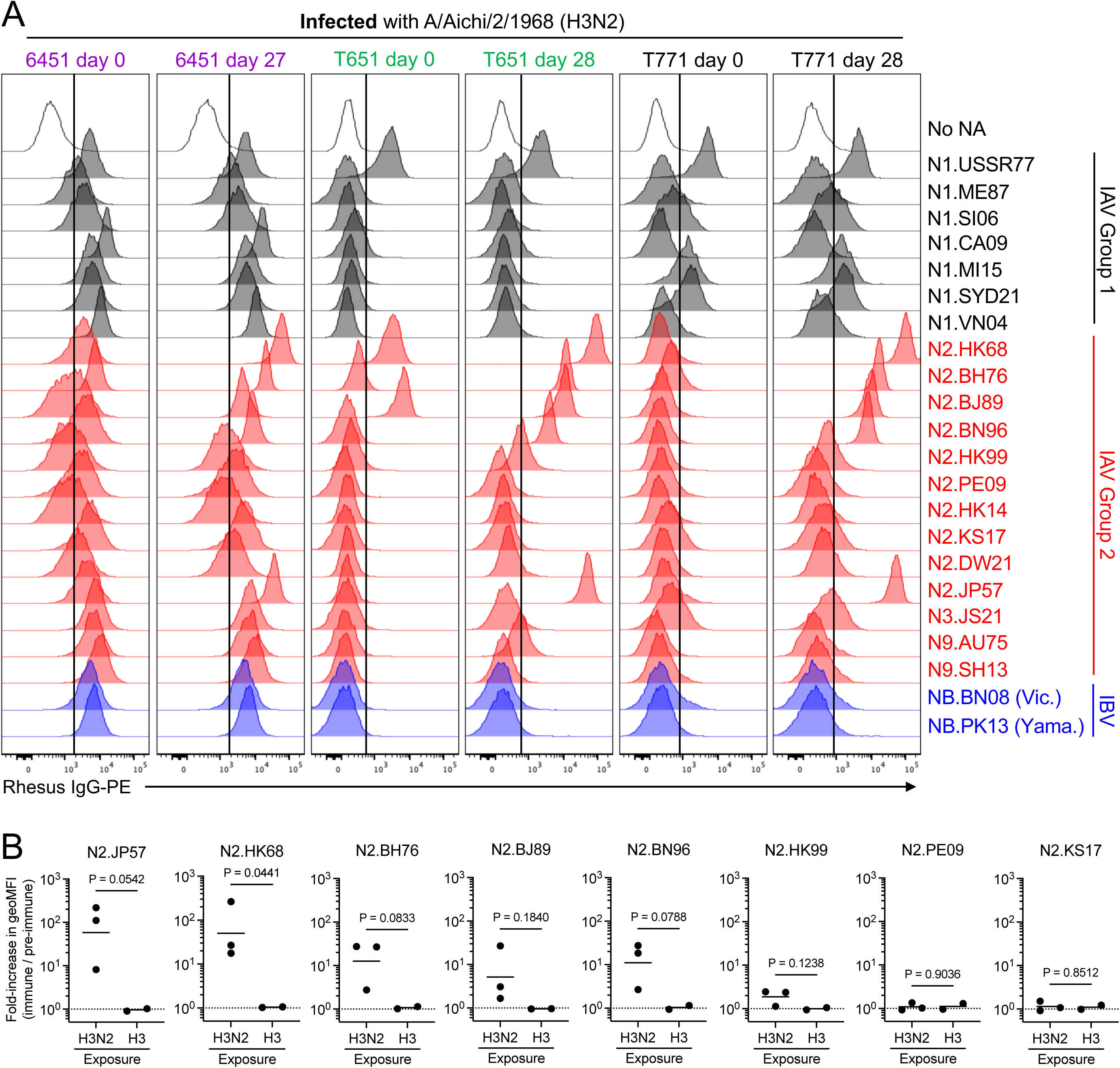
Analysis of serum IgGs elicited by infection of rhesus macaques with H3N2 influenza virus. A) Pre-immune (day 0) and immune (day 27 or 28) blood plasma from three rhesus macaques (6451, T651, T771) infected with A/Aichi/2/1968 (H3N2) influenza virus was incubated with K530-NA cell lines. The degree of IgG labeling of each cell line was determined by flow cytometry, as in Fig. 1. Vertical, black lines denote the threshold for specific labeling of cell-surface NA, determined according to the labeling intensity observed for K530 cells expressing no NA (top row). B) Fold-change in the fluorescence intensity of plasma IgG labeling of selected K530-N2 cell lines, after either infection with H3N2 virus (Fig. 4A) or immunization with recombinant H3 HA (Supplemental Fig. 3B). Fold-change was calculated as the ratio of the geometric mean fluorescence intensity (geoMFI) resulting from labeling with the immune plasma IgG, divided by the geoMFI resulting from labeling with pre-immune plasma IgG. Each symbol represents a single animal. The difference in group means was analyzed by two-tailed *t*-test with Welch’s correction, as described in Materials and Methods.

### K530-NA cell lines have NA catalytic activity

K530-NA cells are compatible with commonly used sialidase activity assays. In the NA-Star assay, which uses a chemiluminescent, small-molecule substrate (27), K530 cell lines expressing N1, N2, N3, N9, or NB each had NA activity ≥100-fold greater than the activity present in control K530 cells expressing no NA (Fig. 5A). Control mAb S1V2-72 did not inhibit the NA activity of these cell lines, but NA catalytic site-binding IgGs 1G01 or 1G05 inhibited the NA activity in a dose-dependent manner, with half-maximal inhibitory concentrations (IC_50_) comparable to reported values (Fig. 5A, Table 2) (13, 15, 18). In contrast, mAbs T3-P10F5, T3-P27B8, and T1-P30E7 did not significantly inhibit the sialidase activity (Fig. 5A) of K530-NA cell lines to which the corresponding mAbs bound tightly (Fig. 3A). We conclude that the latter mAbs do not substantially cover the catalytic pocket of NA. 1G01 and 1G05 had comparable IC_50_ values regardless of whether NA-Star or fluorogenic 4-methylumbelliferyl-N-acetyl-α-*D*-neuraminic acid (MUNANA) (28) was used as the NA substrate (Supplemental Figure 4, Table 2).

**FIG. 5.**
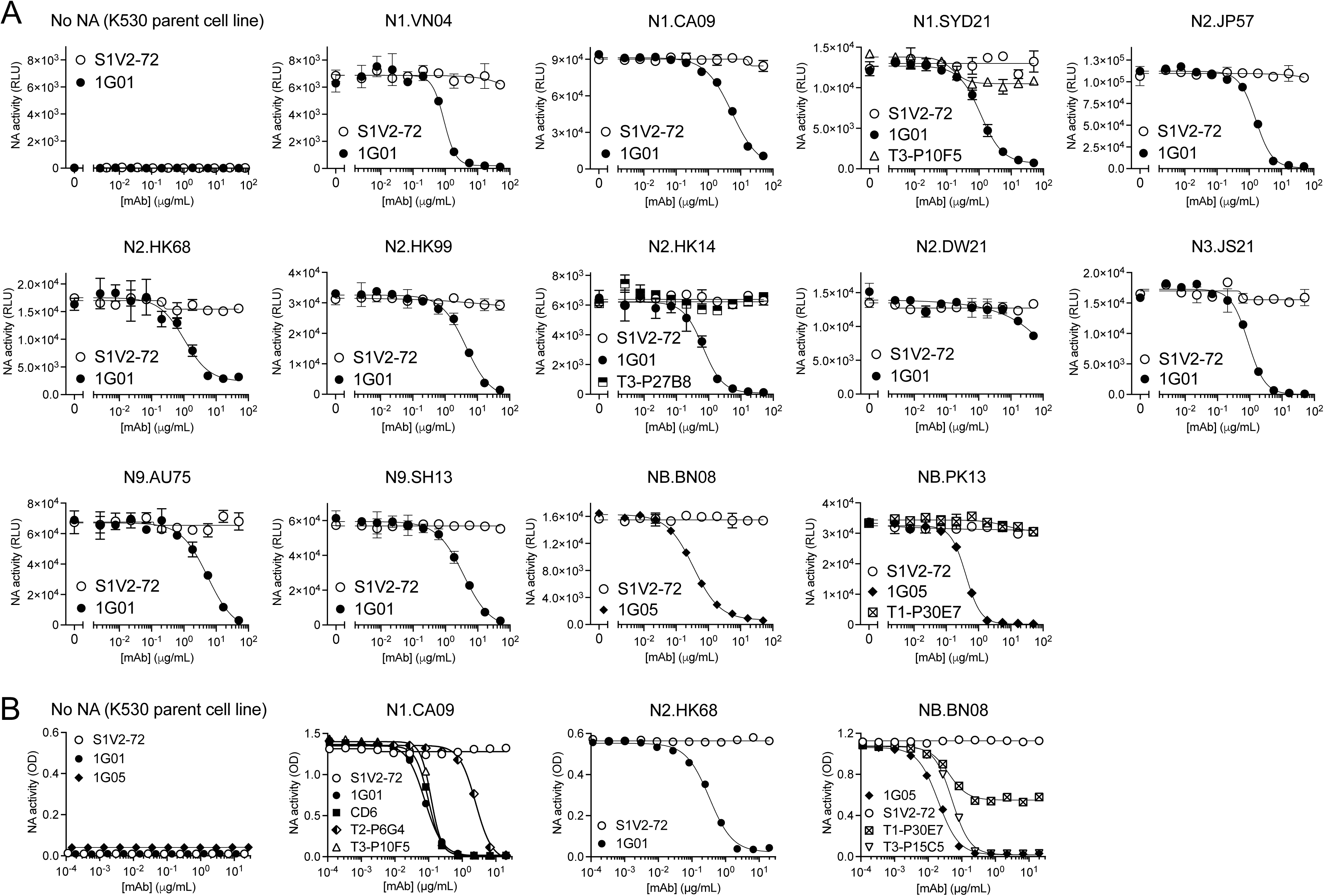
NA catalytic activity of K530-NA cell lines. A) The NA-Star assay was used to determine the sialidase activity of selected K530-NA cell lines in the presence of serially diluted, NA-binding or control rIgGs. B) The enzyme-linked lectin assay (ELLA) was used to measure the sialidase activity of selected K530-NA cell lines in the presence of serially diluted rIgGs. Error bars represent mean ± S.D.

**Table 2.** IC_50_ values for mAb-mediated inhibition of sialidase activity in assays using small-molecule substrates.

| NA strain | 1G01 IC<sub>50</sub> (μg/ml) |  | 1G05 IC<sub>50</sub> (μg/ml) |  | S1V2-72 IC<sub>50</sub> (μg/ml) |  |
| --- | --- | --- | --- | --- | --- | --- |
|  | NA-Star | MUNANA | NA-Star | MUNANA | NA-Star | MUNANA |
| N1.CA09 | 5.4 | 1.4 | - | - | >50 | >50 |
| N1.MI15 | 2.6 | - | - | - | >50 | - |
| N1.SYD21 | 1.1 | - | - | - | >50 | - |
| N1.VN04 | 0.9 | - | - | - | >50 | - |
| N2.JP57 | 1.6 | - | - | - | >50 | - |
| N2.HK68 | 1.0 | 1.1 | - | - | >50 | >50 |
| N2.HK99 | 4.5 | - | - | - | >50 | - |
| N2.HK14 | 0.7 | - | - | - | >50 | - |
| N2.DW21 | >50 | - | - | - | >50 | - |
| N3.JS21 | 0.9 | - | - | - | >50 | - |
| N9.AU75 | 5.5 | - | - | - | >50 | - |
| N9.SH13 | 3.6 | - | - | - | >50 | - |
| NB.BN08 | - | - | 0.3 | 0.4 | >50 | >50 |
| NB.PK13 | - | - | 0.4 | - | >50 | - |

Abs that inhibit NA function without directly blocking the catalytic pocket can be identified by their ability to inhibit cleavage of sialosides from glycosylated protein in the enzyme-linked lectin assay (ELLA) (29). MAbs 1G01 and 1G05 inhibited the sialidase activity of K530-NA cell lines in ELLA, while control mAb S1V2-72 did not (Fig. 5B, Table 3). MAb CD6, which sterically hinders sialoside cleavage (21), inhibited the NA activity of K530-N1.CA09 cells; so did T3-P10F5 and T2-P6G4 (Fig. 5B, Table 3), with IC_50_ values proportional to their relative avidities (as measured by flow cytometry) (Fig. 2). T3-P15C5 potently blocked the activity of K530-NB.BN08 cells (Fig. 5B and Table 3), while T1-P30E7 inhibited NB.BN08 activity by only 50%, even at high IgG concentrations, suggesting partial steric or allosteric inhibition.

**Table 3.** IC_50_ values for mAb-mediated inhibition of sialidase activity in ELLA.

| NA strain | ELLA IC<sub>50</sub> (μg/ml) |  |  |  |  |  |  |  |
| --- | --- | --- | --- | --- | --- | --- | --- | --- |
|  | 1G01 | 1G05 | CD6 | T1-P30E7 | T2-P6G4 | T3-P10F5 | T3-P15C5 | S1V2-72 |
| N1.CA09 | 0.08 | - | 0.10 | - | 2.50 | 0.12 | - | >40 |
| N2.HK68 | 0.33 | - | - | - | - | - | - | >40 |
| NB.BN08 | - | 0.02 | - | 0.52 | - | - | 1.06 | >40 |

## DISCUSSION

Recombinant, soluble NA ectodomains are often challenging to isolate in quantities sufficient for studies of NA Ab responses. A few alternate approaches to NA expression have been described, to which we now add display of native, full-length NAs anchored to the surface of fluorescently barcoded cell lines. These stable cell lines are simple to culture and easy to share among labs. Inclusion of the native transmembrane and stalk domains of NA should enable the NA on K530-NA lines to have the proper antigenic structure (30, 31). That K530-NA cell lines have robust catalytic activity and are avidly and specifically bound by well-characterized, standard mAbs implies that the membrane-bound NAs assume the native, multimeric conformation (13–15, 18, 21, 32).

An advantage of K530-NA cells is their display of full-length NA, i.e., both head and stalk domains. In principle, inclusion of the stalk domain enables discovery of anti-stalk Abs, which have not yet been reported, and which would be invisible in experiments that use only the NA head domain. MAb T3-P20D5 may be one such Ab; alternatively, it may bind some other epitope not represented well on tetrameric, head-only NAs.

The sialic acid-binding activity of K530-NA cell lines might be expected to contribute to non-specific binding of sialylated IgGs, but our results exclude this hypothesis. Pre-immune macaque serum, which presumably contains much sialylated IgG (33, 34), generally labeled NA-expressing cell lines no more intensely than it did control cells that expressed no NA. To the extent that pre-immune serum IgG did bind NA-expressing cell lines, it bound specific NAs rather than all NAs, and did so only in some animals. Therefore, such binding was due to the specificity of the IgG paratope(s), rather than the interaction of IgG-linked sialic acid with NA. Additionally, recombinantly expressed control IgG and ∼1,800 clonal IgGs secreted from cultured human Bmem cells bound NA-expressing and -nonexpressing cell lines with equal intensity. Thus, the sialic acid-binding activity of K530-NA cells does not contribute appreciably to the binding of either recombinant or native IgG.

## ACKNOWLEDGMENT

The research was supported by NIAID grant number P01 AI089618. We thank all members of the consortium participating in this program project for discussions throughout the course of the research reported here. We are grateful for the excellent technical support of Amanda Foreman, Wenli Zhang, Steven Slater, Stephanie H. Smith and Darren J. Morrow at Duke University. This research was conducted in part using equipment and services provided by the Harvard Medical School Immunology Flow Cytometry Core Facility, the Duke University DNA Analysis Facility, and the Duke Human Vaccine Institute Research Flow Cytometry Shared Resource Facility.

SCH is an Investigator in the Howard Hughes Medical Institute. Accession codes for all NA sequences are listed in Supplemental Table 1. We gratefully acknowledge all data contributors, i.e., the Authors and their Originating laboratories responsible for obtaining the specimens, and their Submitting laboratories for generating the genetic sequence and metadata and sharing via the GISAID Initiative (35), on which this research is based. These contributors include the WHO Collaborating Centre for Reference and Research on Influenza (Melbourne, Australia), Queensland Health Forensic and Scientific Services (Coopers Plains, Australia), National Institute for Medical Research (London, United Kingdom), Crick Worldwide Influenza Centre (London, United Kingdom), Erasmus Medical Center (Rotterdam, Netherlands), National Institute for Public Health and the Environment (Bilthoven, Netherlands), New York Medical College (Valhalla, USA), U.S. Centers for Disease Control and Prevention (Atlanta, USA), WHO National Influenza Centre (Nonthaburi, Thailand), Jiangsu Provincial Center for Disease Control & Prevention (Nanjing, China), and the WHO Chinese National Influenza Center (Beijing, China).

## MATERIALS AND METHODS

### Study approvals and volunteers

The study procedures, informed consent, and data collection documents were reviewed and approved by the Duke Health Institutional Review Board (Pro00020561, initial approval 2010). We enrolled three donors (T1, T2, T3) between the ages of 13 yo and 18 yo. Each donor had a documented history of receiving seasonal influenza vaccines ≥3 times prior to the 2019-2020 northern hemisphere flu season. Written informed consent was obtained from each donor’s parent and assent obtained for each participant. Each donor was administered the 2019-2020 quadrivalent inactivated influenza vaccine, and blood was collected 15 d (T1, T2) or 14 d (T3) post-vaccination. Blood samples were processed into PBMCs and plasma and then aliquoted and stored in liquid nitrogen vapor phase or at −80°C, respectively, for future analysis.

### Animal Studies

Five adult rhesus macaques (*Macaca mulatta*) were housed at BioQual and maintained in accordance with the Association for Accreditation of Laboratory Animal Care guidelines at the NIH. Three animals were infected with H3N2 A/Aichi/2/1968 influenza virus at 2.36×10^6^ plaque-forming units per animal in a divided dose: for each animal, half of the infectious dose was given in 1 mL intranasally and the other half as 1 mL intratracheally. Animals were monitored for signs of infection and no intervention was needed. The other two animals were immunized intramuscularly with 100 μg of H3 A/Aichi/2/1968 recombinant protein per animal per time point given as 500-μL total injection volume divided into two sites—250 μL each at right and left quadriceps. The final immunization mixture contained 15% Span85-Tween 80-squalene+R848+CpG oligodeoxynucleotides adjuvant (36) with the remaining volume being sterile saline. Blood was collected in EDTA tubes per the schedule shown in Supplemental Fig. 3A and processed for plasma and cells, which were cryopreserved until use.

### Cell line culture

Unless otherwise noted, mammalian cell lines were maintained in static cultures at 37°C with 5% CO_2_ in a humidified incubator, and culture reagents were from Gibco. The MEC-147 cell line (manuscript in preparation), a derivative of MS40L-low feeder cells (*Mus musculus*)(37, 38) that stably expresses human IL-2, IL-4, IL-21, and BAFF, was expanded from frozen aliquots in Iscove’s Modified Dulbecco’s Medium (IMDM) containing 10% HyClone FBS (Cytiva), 2-mercaptoethanol (55 μM), penicillin (100 units/ml), and streptomycin (100 μg/ml). Lenti-X 293T cells (*Homo sapiens*, Takara) were cultured in Dulbecco’s Modified Eagle Medium (DMEM) plus 10% FBS, penicillin, streptomycin, HEPES (10 mM), sodium pyruvate (1 mM), and 1× MEM non-essential amino acids. Expi293F cells (*Homo sapiens*; Thermo Fisher) were cultured in Expi293 Expression Medium plus penicillin and streptomycin, at 8% CO_2_ with shaking. K530-derived cell lines (*Homo sapiens*) (23) were initially cultured in RPMI-1640 medium plus 10% FBS, 2-mercaptoethanol, penicillin, streptomycin, HEPES, sodium pyruvate, and MEM nonessential amino acids, but IMDM plus 10% FBS, penicillin, and streptomycin was later chosen as the standard growth medium. High Five cells (BTI-TN-5B1-4; *Trichoplusia ni*; Thermo Fisher) were maintained in ESF 921 medium (Expression Systems) at 28°C in spinner flasks in air. Cell lines were not subject to authentication.

### Generation of NA-expressing K530 cell lines

pMD2.G (Addgene plasmid #12259) and psPAX2 (Addgene plasmid #12260) were gifts from Didier Trono. Codon-optimized, NA-encoding DNA sequences were cloned into the pLB-EXIP lentiviral transfer plasmid (23). Influenza strain designations and abbreviated names are shown in Table 1. Accession codes for the amino acid sequences of the corresponding NAs are listed in Supplemental Table 1. Transfer plasmid, pMD2.G and psPAX2 were co-transfected into Lenti-X 293T cells to produce lentivirus. NA-expressing lentiviruses were used to transduce K530 cell lines expressing unique combinations of EBFP2, mTurquoise2, mNeonGreen, and mCardinal (23). Transduced K530 cells were cultured for one week, then labeled with 1G01 (18) or DA03E17 (15) human IgG1 (2 μg/ml), washed, and labeled with goat anti-human IgG-PE (Southern Biotech; 2 μg/ml). Individual cells expressing high levels of NA were identified by flow cytometry and then sorted into 96-well plates containing growth medium. Subclones were expanded for ∼10 days, then re-analyzed by flow cytometry to identify subclones with uniform, high expression of NA. For each cell line, one rapidly growing subclone with high NA expression was selected for use in all subsequent experiments. Aside from selecting subclones with high NA expression, no effort was made to select cell lines with comparable expression levels of NA. For long-term storage, K530 cell lines suspended in 90% FBS plus 10% DMSO and cryo-preserved in liquid nitrogen.

### Expression and purification of recombinant, tetrameric NA heads

Recombinant NAs were soluble, head-only, tetrameric ectodomains (20). NAs were expressed by infection of insect cells with recombinant baculovirus as described (20, 39–42). In brief, a pFastBac vector was modified to encode a secretion signal peptide, an N-terminal His_8_ tag, an HRV3C protease cleavage site, a tetrabrachion (*Staphylothermus marinus*) tetramerization tag, a thrombin cleave site, and the globular head domain of NA. The resulting baculoviruses produce tetrameric NA heads. Supernatant from recombinant baculovirus-infected High Five cells was harvested 72 h post-infection and clarified by centrifugation. Proteins were purified by adsorption to cobalt-nitrilotriacetic acid (Co-NTA) agarose resin (Takara), followed by a wash in buffer A (10 mM Tris, 150 mM NaCl, pH 7.5) plus 5 mM imidazole, elution in buffer A plus 350 mM imidazole (pH 8), and gel filtration chromatography on a Superdex 200 column (GE Healthcare) in buffer A.

### Bmem sorting and culture

PBMCs in RPMI medium plus 10% FBS were incubated with irrelevant mouse IgG1 (MG1K; Rockland) to block nonspecific binding and then labeled with fluorochrome-conjugated mAbs. The following human surface antigen-specific mAbs, purchased from BD Biosciences, BioLegend, or Thermo Scientific, were used: anti-human IgM-fluorescein isothiocyanate (FITC) (MHM-88), anti-CD3-PE-Cy5 (UCHT1), anti-CD14-Tri (TuK4), anti-CD16-PE-Cy5 (3G8), anti-CD19-PE-Cy7 (HIB19), anti-IgG-allophycocyanin (APC) (G18-145), anti-IgD-PE (IA6-2), anti-CD27-BV421 (M-T271), and anti-CD24-BV510 (ML5) Abs. Labeled cells were sorted using a FACSAria II with Diva software (BD Biosciences). Flow cytometric data were analyzed with FlowJo software (Tree Star, Inc.). Total Bmem (CD19^+^ Dump^−^ CD27^+^ CD24^+^) (Supp. Fig. 1A) were identified as described (25, 38, 43). Surface IgD, IgM, and IgG expression were also determined, but were not considered for sorting. Doublets were excluded from cell sorting by forward scatter area (FSC-A) versus FSC height (FSC-H) gating. Cells positive for 7-aminoactinomycin D (7-AAD) (BD Bioscience) or for CD3, CD14, or CD16 expression were also excluded.

Sorted single Bmem were expanded in the presence of feeder cells as described (25, 38, 43), with some modifications. MEC-147 feeder cells (manuscript in preparation) were used instead of MS40L-low feeder cells and exogenous cytokines. Single Bmem were sorted directly into 96-well plates containing feeder cells and 200 μl growth medium per well. After seven days of co-culture, 100 μl of spent medium was removed from each well and replaced with 200 μl of fresh growth medium. On culture days ∼14, ∼17, and ∼21, two-thirds of the spent medium from each well was replaced with an equal volume of fresh growth medium. On culture day 25, culture supernatants were harvested to screen the secreted clonal IgGs. Expanded clonal B cells were frozen at −80°C for V(D)J sequence analysis.

### Luminex multiplex binding assay

The specificities and avidities of clonal IgGs in culture supernatants were determined in a multiplex bead assay (Luminex Corp.) as described (25, 44) with modifications. Culture supernatants were diluted in Luminex assay buffer (phosphate-buffered saline [PBS] containing 1% bovine serum albumin [BSA], 0.05% NaN_3_, and 0.05% Tween 20) with 1% milk and incubated for 2_h with the mixture of antigen-coupled microsphere beads in 96-well filter-bottom plates (Millipore). After washing three times with assay buffer, the beads were incubated for 1_h with PE-conjugated mouse anti-human IgG Abs (JDC-10; Southern Biotech). After three washes, the beads were resuspended in assay buffer and the plates read on a Bio-Plex 3D suspension array system (Bio-Rad). Antigens and controls included BSA, mouse anti-human Ig(κ) (SB81a; Southern Biotech), mouse anti-human Ig(λ) (JDC-12; Southern Biotech), mouse anti-human IgG (Jackson ImmunoResearch), tetanus toxoid from *Clostridium tetani* (List Biological Laboratories), keyhole limpet hemocyanin (KLH; Sigma), ovalbumin (OVA; Sigma), insulin (Sigma), and a panel of recombinant, tetrameric, head-only neuraminidase constructs representing N1 A/California/07/2009; N1 A/Michigan/45/2015, X-275; N2 A/Aichi/2/1968; N2 A/Hong Kong/4801/2014, X263B; B/Phuket/3073/2013; B/Brisbane/60/2008.

### Ab *V(D)J* rearrangement amplification and analysis

Rearranged V(D)J gene sequences were obtained from cultures of clonally expanded human Bmem as described (45). V(D)J rearrangements were identified with Cloanalyst (46) and IMGT/V-QUEST (47).

### Recombinant IgG expression and purification

AbVec2.0-IGHG1, AbVec1.1-IGKC, and AbVec2.1-IGLC2-MscI plasmids, which harbor the constant regions of human IgG1, Igκ, or Igλ, were gifts from Hedda Wardemann (RRID:Addgene_80795; RRID:Addgene_80796; RRID:Addgene_80797)(48). Synthetic DNAs encoding Ab heavy-or light-chain variable domains were cloned into these expression vectors. The vectors were transiently transfected into Expi293F cells with the Expifectamine 293 transfection kit (Thermo Fisher), according to the manufacturer’s instructions. Five days post-transfection, supernatants were harvested, clarified by low-speed centrifugation, mixed 1:1 with Protein A binding buffer, and incubated overnight with Pierce Protein A agarose resin (Thermo Fisher). The resin was collected in a chromatography column, washed with binding buffer, eluted in Pierce IgG Elution Buffer (Thermo Fisher), neutralized by one-tenth volume of 1M Tris (pH 9), and dialyzed into PBS plus 0.1% sodium azide. IgG concentrations were determined with a NanoDrop spectrophotometer (Thermo Fisher).

### Flow cytometry analysis of rIgG binding to K530 cell lines

Pooled K530-NA cell lines were thawed from cryopreserved aliquots and expanded in culture for ≥3 days. Pooled K530-NA cells were incubated at room temperature (RT) for 25-30 min with 2 μg/ml rIgGs diluted in IMDM plus 10% FBS. Alternately, cells were incubated with culture supernatants containing clonal human IgG (diluted 1:20) or with rhesus macaque plasma (diluted 1:40). After washing, cells were labeled with 2 μg/ml PE-conjugated goat anti-human IgG (Southern Biotech) or 2 μg/ml PE-conjugated mouse anti-monkey IgG (SB108a, Southern Biotech) for 20-30 min at RT. Cells were then washed, stained with propidium iodide to identify dead cells, and analyzed with a BD FACSymphony A1 flow cytometer.

### NA-Star assay

NA activity was determined with the NA-Star Influenza Neuraminidase Inhibitor Resistance Detection Kit (Invitrogen). K530-NA cells were suspended in calcium-containing assay buffer (Hank’s Balanced Salt Solution [Gibco] plus 0.5% BSA) and dispensed (1×10^5^ cells/25 μl/well) into opaque, white, 96-well microtiter plates (Thermo Fisher). Recombinant IgGs were serially diluted in assay buffer and added (25 μl/well) to the cells, mixed, and incubated at RT for 30 min. NA-Star substrate was diluted 1:1,000 in assay buffer, added to each well (10 μl/well), and incubated with the cell-Ab mix for 20 min at RT. After adding NA-Star Accelerator solution (60 μl/well), chemiluminescence was detected immediately with a Tecan Spark plate reader, using an integration time of 1 s per well. Inhibition of NA activity was calculated as the percentage of residual NA activity relative to wells containing no inhibitor.

### 4-methylumbelliferyl-N-acetyl-α-*D*-neuraminic acid (MUNANA) cleavage assay

Established protocols (28, 49) were adapted to use K530-NA cell lines as the source of NA. K530-NA cells were suspended in calcium-containing assay buffer (Hank’s Balanced Salt Solution [Gibco] plus 0.5% BSA) and dispensed in 50 μl aliquots into opaque, black, 96-well microtiter plates (Corning) at 1.2×10^3^ cells/well (K530-N1.CA09), 2×10^5^ cells/well (K530-N2.HK68), or 4×10^4^ cells/well (K530-NB.BN08). Recombinant IgGs were serially diluted in assay buffer and added (25 μl/well) to the cells, mixed, and incubated at RT for 20 min.

MUNANA (Biosynth) was diluted in assay buffer and then added to each well (25 μl/well, 0.15 mM final), and then the plate was sealed with adhesive film and was incubated for 60 min at 37°C with gentle shaking (200 rpm). Sialidase activity was quenched with 100 μl/well of 140 mM NaOH plus 83% (v/v) ethanol, and fluorescence intensity was measured with a Tecan Spark plate reader, using an excitation wavelength of 355 nm and an emission wavelength of 460 nm.

### Enzyme-linked lectin assay (ELLA)

High-binding, half-area, 96-well microplates (Greiner Bio-One) were coated with fetuin (MilliporeSigma) diluted to 25 μg/ml in 0.1 M sodium carbonate-bicarbonate buffer (pH 9). Plates were coated with fetuin for 2-3 h at RT, then washed extensively with PBS plus 0.1% Tween-20 (PBS-T). Recombinant IgGs were serially diluted in ELLA assay buffer (IMDM plus 0.5% BSA, 0.5% Tween-20, and penicillin-streptomycin), then added to the microplate (25 μl/well).

K530-NA cells suspended in ELLA assay buffer were added to each well (25 μl/well; K530-N1.CA09, 10^3^ cells/well; K530-N2.HK68, 2×10^5^ cells/well; K530-NB.BN08, 10^5^ cells/well) and mixed with the IgGs by pipetting. The plates were loosely covered and incubated at 37°C, 5% CO_2_ in a humidified environment for 16 h. After extensive washing with PBS-T, horseradish peroxidase-conjugated peanut agglutinin (MilliporeSigma) diluted in PBS plus 0.5% BSA was added to each well and incubated for 2 h at RT. After washing with PBS-T, peroxidase activity was detected with the TMB substrate kit (BioLegend) and a Tecan Spark plate reader. Background signal at 650 nm was subtracted from the signal at 450 nm to calculate the OD450.

### Statistical analysis

Geometric mean fluorescence intensity values were log-transformed, and differences between means of the infected and immunized groups were calculated by an unpaired t test (two-tailed) with Welch’s correction for unequal variances, using GraphPad Prism v10.4.2. Differences were considered statistically significant at P < 0.05.

## Abbreviations

Ab: antibody Bmem memory B cell
IAV: influenza A virus
IBV: influenza B virus
HA: hemagglutinin
NA: neuraminidase
rIgG: recombinant IgG
BSA: bovine serum albumin
KLH: keyhole limpet hemocyanin
OVA: chicken ovalbumin
RT: room temperature
TT: tetanus toxoid
mAb: monoclonal antibody

## Data availability

V(D)J sequences for mAbs from donors T1, T2, and T3 are available at GenBank (www.ncbi.nlm.nih.gov/Genbank), accession numbers PQ818731-PQ818752.

## CRediT Statement

**Joel Finney**: Conceptualization, Investigation, Visualization, Writing – Original Draft, Writing – Review and Editing; **Masayuki Kuraoka**: Investigation, Writing – Review and Editing; **Shengli Song**: Resources, Writing – Review and Editing; **Akiko Watanabe**: Investigation**; Xiaoe Liang**: Investigation**; Dongmei Liao**: Investigation**; M. Anthony Moody**: Resources, Writing – Review and Editing; Funding Acquisition**; Emmanuel B. Walter**: Resources**; Stephen C. Harrison**: Writing – Review and Editing; Funding Acquisition**; Garnett Kelsoe**: Conceptualization, Supervision, Writing – Review and Editing; Funding Acquisition

## ICMJE criteria for authorship

All authors attest they meet the ICMJE criteria for authorship.

