## Supplemental Figures for "Fluorescence-barcoded cell lines stably expressing membrane-anchored influenza neuraminidases"

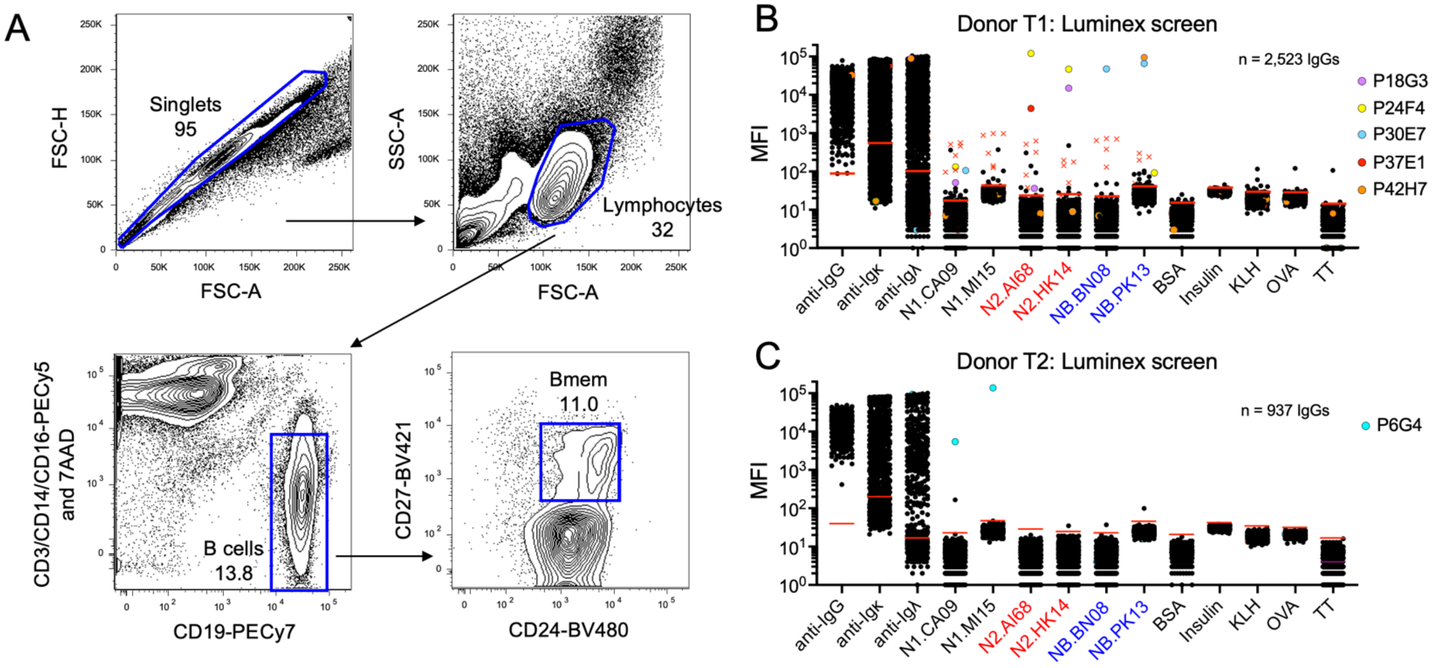


**Supplemental Figure 1. Isolation of IgGs from human memory B (Bmem) cells.** A) Flow cytometry plots showing the sorting strategy for isolation of Bmem. The plots are from donor T1; the same strategy was used for sorting from other donors' samples. Inset labels denote the identity of the gated population and its frequency within the parent population. B-C) Luminex assay results showing the antigen-binding profiles of clonal IgG-containing culture supernatants from Bmem cells isolated from donors T1 (B) or T2 (C). Short, red, horizontal lines in each column denote the limit of detection, calculated as six standard deviations above the mean signal produced by cultures containing no B cells. Clonal IgGs of interest are denoted with uniquely colored symbols. IgGs denoted by red "X"s in (B) correspond to samples for which binding to influenza hemagglutinin (HA) was ≥100-fold higher than binding to NA (data not shown). These samples were considered to be HA-specific IgGs rather than NA-specific IgGs.

A

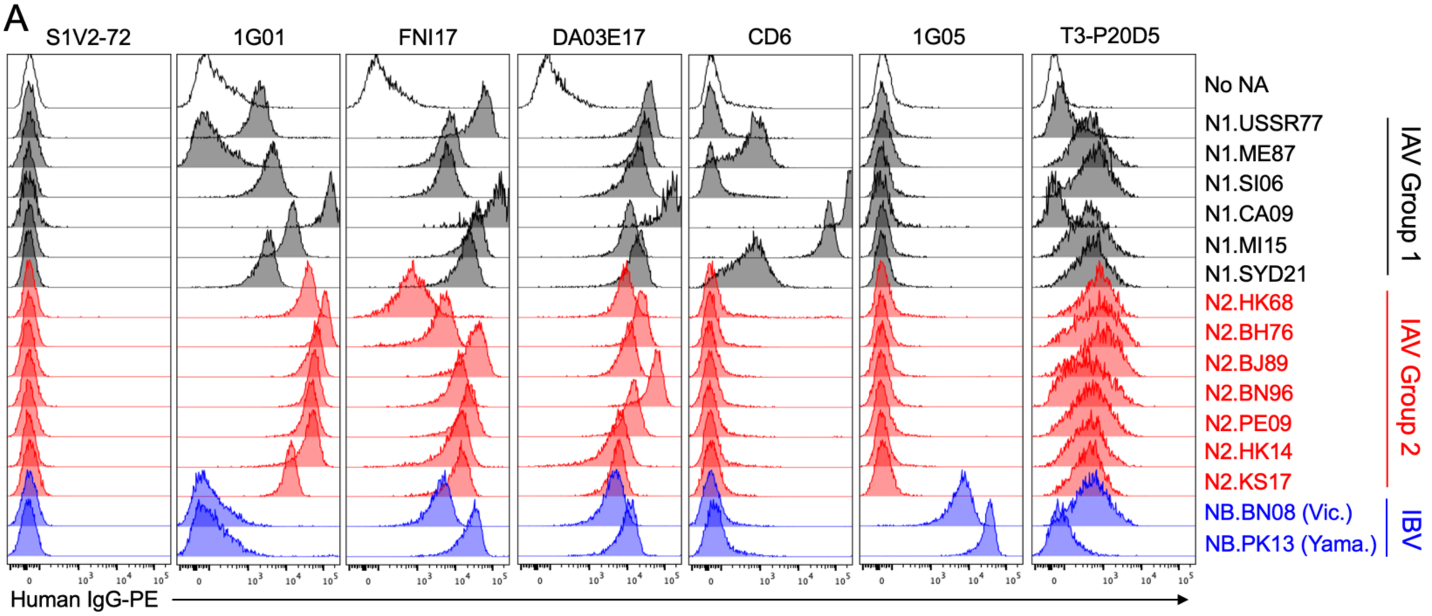


B Luminex MFIs:

| | Anti-IgG | N1 CA09 | N1 MI15 | N2 AI68 | N2 HK14 | NB BN08 | NB PK13 |
|---|---|---|---|---|---|---|---|
| T3-P20D5 | 23,749 | 6 | 33 | 7 | 16 | 9 | 38 |

| |
|---|
| ≥ 10,000 |
| ≥ 1,000 |
| < 1,000 |
| < LOD |

**Supplemental Fig. 2. MAb T3-P20D5 specifically labels K530-NA cells.** A) Shown are flow cytometry histograms depicting the binding of recombinant IgG versions of NA mAbs to K530 cell lines expressing membrane-anchored NAs, as in Fig 1. MAbs were incubated with pooled cell lines comprising Option 1. The detector for phycoerythrin (PE) fluorescence, used for measuring IgG binding, was set at a higher voltage than for a typical experiment. As a result, the signal for some samples (e.g. CD6 binding to N1.CA09) exceeds the detector's upper limit, while the signal for T3-P20D5 binding to certain NAs is clearly distinguishable from its lack of binding to the control K530 cell line that expresses no NA. B) Luminex binding assay results for the IgG-containing culture supernatant from sample T3-P20D5. The sample contains a significant quantity of IgG, but binding to recombinant, tetrameric NA heads is below the limit of detection (LOD; calculated as in Fig. 2A).

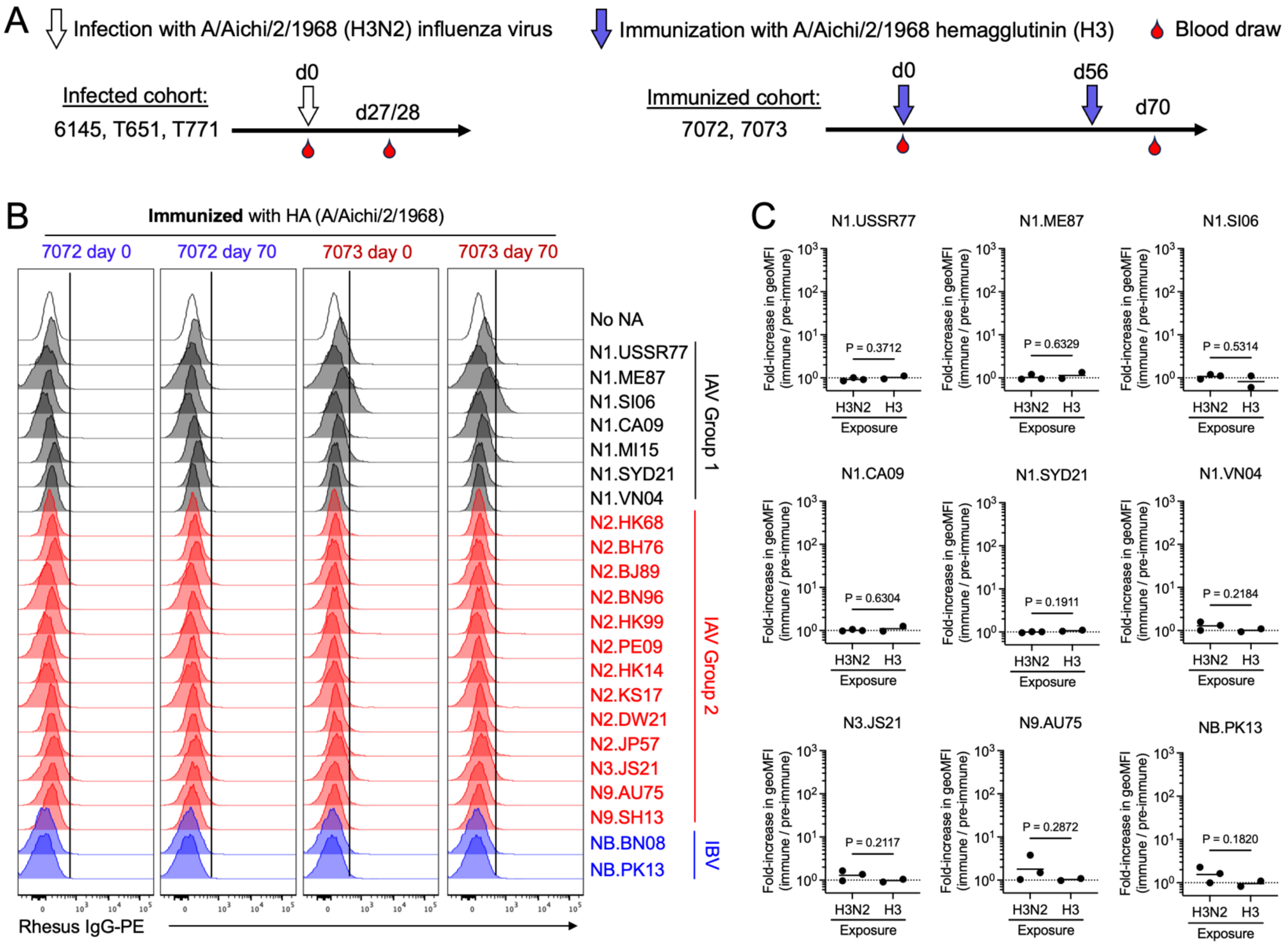


**Supplemental Figure 3. Analysis of the serum IgG response to influenza infection in rhesus macaques.** A) The timing of infection (empty downward arrows), immunizations (blue downward arrows), and blood draws (red teardrops) are shown along a timeline. The days, denoted with a "d," beginning on the day of immunization or infection (d0), are indicated above each line. The animals in each cohort are indicated. B) Pre-immune (day 0) or immune (day 70) blood plasma from two rhesus macaques (7072, 7073) immunized twice with A/Aichi/2/1968 (H3) hemagglutinin was incubated with K530 cell lines expressing membrane-anchored NAs. The degree of IgG labeling of each cell line was determined by flow cytometry, as in Fig. 1. Vertical, black lines denote the threshold for specific labeling of cell-surface NA, determined according to the labeling intensity observed for K530 cells expressing no NA (top row). C) Fold-change in the fluorescence intensity of plasma IgG labeling of selected K530-NA cell lines, after either infection with H3N2 virus (Fig. 4A) or immunization with recombinant H3 HA (Supplemental Fig. 3B). Fold-change was calculated as the ratio of the geometric mean fluorescence intensity (geoMFI) resulting from labeling with the immune plasma IgG, divided by the geoMFI resulting from labeling with pre-immune plasma IgG. Each symbol represents a single animal. The difference in group means was analyzed by two-tailed *t*-test with Welch's correction, as described in Materials and Methods.

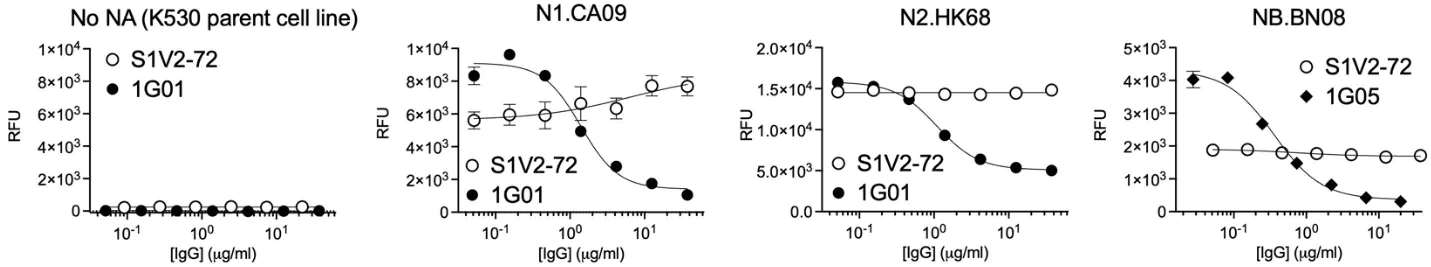


**Supplemental Figure 4. NA catalytic activity of K530-NA cell lines in a MUNANA cleavage assay.** The sialidase activity of selected K530-NA cell lines was determined in the presence of serially diluted, catalytic site-binding or control rIgGs. Error bars represent mean ± S.D of three replicates.
